# A comprehensive evaluation of long read error correction methods

**DOI:** 10.1101/519330

**Authors:** Haowen Zhang, Chirag Jain, Srinivas Aluru

## Abstract

**Background:** Third-generation single molecule sequencing technologies can sequence long reads, which is advancing the frontiers of genomics research. However, their high error rates prohibit accurate and efficient downstream analysis. This difficulty has motivated the development of many long read error correction tools, which tackle this problem through sampling redundancy and/or leveraging accurate short reads of the same biological samples. Existing studies to asses these tools use simulated data sets, and are not sufficiently comprehensive in the range of software covered or diversity of evaluation measures used.

**Results:** In this paper, we present a categorization and review of long read error correction methods, and provide a comprehensive evaluation of the corresponding long read error correction tools. Leveraging recent real sequencing data, we establish benchmark data sets and set up evaluation criteria for a comparative assessment which includes quality of error correction as well as run-time and memory usage. We study how trimming and long read sequencing depth affect error correction in terms of length distribution and genome coverage post-correction, and the impact of error correction performance on an important application of long reads, genome assembly. We provide guidelines for practitioners for choosing among the available error correction tools and identify directions for future research.

**Conclusions:** Despite the high error rate of long reads, the state-of-the-art correction tools can achieve high correction quality. When short reads are available, the best hybrid methods outperform non-hybrid methods in terms of correction quality and computing resource usage. When choosing tools for use, practitioners are suggested to be careful with a few correction tools that discard reads, and check the effect of error correction tools on downstream analysis. Our evaluation code is available as open-source at https://github.com/haowenz/LRECE.

## Background

Third-generation sequencing technologies produce long reads with average length of 10 Kbp or more that are orders of magnitudes longer than the short reads available through second-generation sequencing technologies (typically a few hundred bp). In fact, the longest read length reported to date is > 1 million bp [1]. Longer lengths are attractive because they enable disambiguation of repetitive regions in a genome or a set of genomes. The impact of this valuable long-range information has already been demonstrated for *de novo* genome assembly [2, 3, 4], novel variant detection [5, 6], RNA-seq analysis [7], metagenomics [8], and epigenetics [9, 10].

The benefit of longer read lengths, however, comes with the major challenge of handling high error rates. Currently, there are two widely used third-generation single molecule sequencing platforms – Pacific Biosciences (PacBio) and Oxford Nanopore Technologies (ONT). Both sequencing platforms are similar in terms of their high error rates (ranging from 10-20%) with most errors occurring due to insertions or deletions (indels); however the error distribution varies [4, 11, 12]. Pacbio sequencing errors appear to be randomly distributed over the sequence [13]. For ONT on the other hand, the error profile has been reported to be biased. For example, A to T and T to A substitutions are less frequent than other substitutions, and indels tend to occur in homopolymer regions [12, 14]. These error characteristics pose a challenge for long read data analyses, particularly for detecting correct read overlaps during genome assembly and variants at single base pair resolution, thus motivating the development of error correction methods.

Error correction algorithms are designed to identify and fix or remove sequencing errors, thereby benefiting resequencing or *de novo* sequencing analysis. In addition, the algorithms should be computationally efficient to handle increasing volumes of sequencing data, particularly in the case of large, complex genomes. Numerous error correction methodologies and software have been developed for short reads; we refer readers to [15] and [16] for a thorough review. Given the distinct characteristics of long reads, i.e., significantly higher error rates and lengths, specialized algorithms are needed to correct them. Till date, several error correction tools for long reads have been developed including PacBioToCA [17], LSC [18], ECTools [19], LoRDEC [20], proovread [21], NaS [22], Nanocorr [23], Jabba [24], CoLoRMap [25], LoRMA [26], HALC [27], FLAS [28], FMLRC [29], HG-CoLoR [30] and Hercules [31].

In addition, error correction modules have been developed as part of long read *de novo* assembly pipelines, such as Canu [32] and HGAP [33]. In the assembly pipeline, correction helps by increasing alignment identities of overlapping reads, which facilitates overlap detection and improves assembly. Many long read error correction tools require and make use of highly accurate short reads to correct long reads (accordingly referred to as hybrid methods). Others, referred to as non-hybrid methods, perform self-correction of long reads using overlap information among them.

A few review studies have showcased comparisons among rapidly evolving error correction algorithms to assess state-of-the-art. Laehnemann *et al.* [34] provide an introduction to error rates/profiles and a methodology overview of a few correction tools for various short and long read sequencing platforms, although no benchmark is included. A review and benchmark for hybrid methods is also available [35]. However, the study only used simulated reads and focused more on speed rather than correction accuracy. Besides, it does not include non-hybrid methods in the assessment. More recently, LRCstats [36] was developed for evaluation of long read error correction software; however, it is restricted to benchmarking with simulated reads.

While benchmarking with simulated reads is useful, it fails to convey performance in real-world scenarios. Besides the base-level errors (i.e., indels and substitutions), real sequencing data sets also contain larger structural errors, e.g., chimeras [37]. However, state-of-the-art simulators (e.g., SimLoRD [38]) only generate reads with base-level errors rather than structural errors. Furthermore, Miclotte *et al.* [24] consistently observed worse performance when using real reads instead of simulated reads, suggesting that simulation may fail to match the characteristics of actual error distribution. Therefore, benchmarking with real data sets is important.

In this study, we establish benchmark data sets, present an evaluation methodology suitable to long reads, and carry out comprehensive evaluation of the quality and computational resource requirements of state-of-the-art long read correction software. We also study the effect of trimming and different sequencing depths on correction quality. To understand impact of error correction on downstream analysis, we perform assembly using corrected reads generated by various tools and assess quality of the resulting assemblies.

## Overview of long read error correction methods

### Hybrid methods

Hybrid methods take advantage of high accuracy of short reads (error rates often < 1%) for correcting errors in long reads. An obvious requirement is that the same biological sample must be sequenced using both short read and long read technologies. Based on how these methods make use of short reads, we further divide them into two categories: *alignment-based* and *assembly-based*. The first category includes Hercules, CoLoRMap, Nanocorr, Nas, proovread, LSC and PacBioToCA, whereas HG-CoLoR, HALC, Jabba, LoRDEC, and ECTools are in the latter. The ideas underlying the methods are summarized below.

#### Short-read-alignment-based methods

As a first step, these methods align short reads to long reads using a variety of aligners, e.g. BLAST [39], Novoalign (http://www.novocraft.com/products/novoalign/). As long reads are usually error-prone, some alignments can be missed or biased. Thus, most of the tools in this category utilize various approaches to increase accuracy of alignments. Drawing upon the alignments, these methods use distinct approaches to generate corrected reads.

##### PacBioToCA

Consensus sequences for long reads are generated by multiple sequence alignment of short reads using AMOS consensus module [40].

##### LSC

Short reads and long reads are compressed using homopolymer compression (HC) transformation prior to alignment. Then error correction is performed at HC points, mismatches and indels by temporarily decompressing the aligned short reads and then generating consensus sequences. Finally, the corrected sequences are decompressed.

##### Proovread

Similar to PacBioToCA and LSC, short reads are mapped to long reads, and the resulting alignments are used to call consensus. But its alignment parameters are carefully selected and adapted to the PacBio sequencing error profile. To further improve correction, the phred quality score and Shannon entropy value are calculated at each nucleotide for quality control and chimera detection, respectively. To reduce run time, an iterative correction strategy is employed. Three pre-correction steps are performed using increasing subsamples of short reads. In each step, the long read regions are masked to reduce alignment search space once they are corrected and covered by a sufficient number of short read alignments. In the final step, all short reads are mapped to the unmasked regions to make corrections.

##### NaS

Like the other tools in this category, it first aligns short reads with long reads. However, only the stringently aligned short reads are found and kept as seed-reads. Then instead of calling consensus, similar short reads are retrieved with these seed-reads. Micro-assemblies of these short reads are performed to generate contigs, which are regarded as corrected reads. In other words, the long reads are only used as template to select seed-reads.

##### Nanocorr

It follows the same general approach as PacBioToCA and LSC, by aligning short reads to long reads and then calling consensus. But before the consensus step, a dynamic programming algorithm is utilized to select an optimal set of short read alignments that span each long read.

##### CoLoRMap

CoLoRMap does not directly call consensus. Instead, for each long read region, it runs a shortest path algorithm to construct a sequence of overlapping short reads aligned to that region with minimum edit distance. Subsequently, the region is corrected by the constructed sequence. In addition, for each uncovered region (called gap) on long reads, any unmapped reads with corresponding mapped mates are retrieved and assembled locally to fill the gap.

##### Hercules

It first aligns short reads to long reads. Then unlike other tools, Hercules uses a machine learning-based algorithm. It creates a profile Hidden Markov Model (pHMM) template for each long read and then learns posterior transition and emission probabilities. Finally, the pHMM is decoded to get the corrected reads.

#### Short-read-assembly-based methods

These methods first perform assembly with short reads, e.g., generate contigs using an existent assembler, or only build the de Bruijn graph (DBG) based on them. Then the long reads are aligned to the assemblies, i.e., contigs/unitigs or a path in the DBG, and corrected. Algorithms for different tools in this category are summarized below.

##### ECTools

First, unitigs are generated from short reads using any available assembler and aligned to long reads. Afterwards, the alignments are filtered to select a set of unitigs which provide the best cover for each long read. Finally, differences in bases between each long read and its corresponding unitigs are identified and corrected.

##### LoRDEC

Unlike ECTools which generates assemblies, LoRDEC only builds a DBG of short reads. Subsequently, it traverses paths in the DBG to correct erroneous regions within each long read. The regions are replaced by the respective optimal paths which are regarded as the corrected sequence.

##### Jabba

It adopts a similar strategy as in LoRDEC, and builds a DBG of short reads followed by aligning long reads to the graph to correct them. The improvement is that Jabba employs a seed-and-extend strategy using maximal exact matches (MEMs) as seeds to accelerate the alignment.

##### HALC

Similar to ECTools, short reads are used to generate contigs as the first step. Unlike other methods which try to avoid ambiguous alignments [17, 41], HALC aligns long reads to the contigs with a relatively low identity requirement, thus allowing long reads to align with their similar repeats which might not be their true genomic origin. Then long reads and contigs are split according to the alignments so that every aligned region on read has its corresponding aligned contig region. A contig graph is constructed with the aligned contig regions as vertices. A weighted edge is added between two vertices if there are adjacent aligned long read regions supporting it. The more regions support the edge, the lower is the weight assigned to it. Each long read is corrected by the path with minimum total weight in the graph. Furthermore, the corrected long read regions are refined by running LoRDEC, if they are aligned to similar repeats.

##### FMLRC

This software uses a DBG-based correction strategy similar to LoRDEC. However, the key difference in the algorithm is that it makes two passes of correction using DBGs with different *k*-mer sizes. The first pass does the majority of correction, while the second pass with a longer *k*-mer size corrects repetitive regions in the long reads. Note that a straightforward implementation of a DBG does not support dynamic adjustment of *k*-mer size. As a result, FMLRC uses FM-index to implicitly represent DBGs of arbitrary length *k*-mers.

##### HG-CoLoR

Similar to FMLRC, it avoids using a fixed *k*-mer size for the de Bruijn graph. Accordingly, it relies on a variable-order de Bruijn graph structure [42]. It also uses a seed-and-extend approach to align long reads to the graph. However, the seeds are found by aligning short reads to long reads rather than directly selecting them from the long reads.

### Non-hybrid methods

These methods perform self-correction with long reads alone. They all contain a step to generate consensus sequences using overlap information. However, the respective methods vary in how they find the overlaps and generate consensus sequences. The details are as follows.

#### FLAS

It takes all-to-all long read overlaps computed using MECAT [43] as input, and clusters the reads that are aligned with each other. In case of ambiguous instances, i.e., the clusters that share the same reads, FLAS evaluates the overlaps by computing alignments using sensitive alignment parameters either to augment the clusters or discard the incor-rect overlaps. The refined alignments are then used to correct the reads. To achieve better accuracy, it also corrects errors in the uncorrected regions of the long reads. Accordingly, it constructs a string graph using the corrected regions of long reads, and aligns the uncorrected ones to the graph for further correction.

#### LoRMA

By gradually increasing the *k*-mer size, LoRMA iteratively constructs DBGs using *k*-mers from long reads exceeding a specified frequency threshold, and runs LoRDEC to correct errors based on the respective DBGs. After that, a set of reads similar to each read termed *friends* are selected using the final DBG, which should be more accurate due to several rounds of corrections. Then, each read is corrected by the consensus sequence generated by its friends.

##### Canu error correction module

As a first step during the correction process, Canu computes all-versus-all overlap information among the reads using a modified version of MHAP [44]. It uses a filtering mechanism during the correction to favor true overlaps over the false ones that occur due to repetitive segments in genomes. The filtering heuristic ensures that each read contributes to correction of no more than *D* other reads, where *D* is the expected sequencing depth. Finally, a consensus sequence is generated for each read using its best set of overlaps by leveraging “falcon sense” consensus module [3].

## Methods

We selected data sets from recent publicly accessible genome sequencing experiments. For benchmarking the different programs, our experiments used genome sequences from multiple species and different sequencing platforms with recent chemistry, e.g., R9 for ONT or P6-C4/P5-C3 for PacBio. We describe our evaluation criteria and use it for a comprehensive assessment of the correction methods/software.

### Benchmark data sets

Our benchmark includes resequencing data from three reference genomes – *Escherichia coli* K-12 MG1655 (*E. coli*), *Saccharomyces cerevisiae* S288C (yeast), and *Drosophila melanogaster* ISO1 (fruit fly). The biggest hurdle when benchmarking with real data is the absence of ground truth (i.e., perfectly corrected reads). However, the availability of reference genomes of these strains enables us to evaluate the output of correction software in a reliable manner using the reference. Essentially, differences in a corrected read with respect to the reference imply uncorrected errors. A summary of the selected read data sets is listed in Table 1. We leveraged publicly available high coverage read data sets of the selected genomes available from all three platforms – Illumina (for short reads), Pacbio, and ONT. In addition, some of these samples were sequenced using multiple protocols, yielding reads of varying quality. This enabled us to do a thorough comparison among error correction software across various error rates and error profiles.

**Table 1.** Details of the benchmark data sets.

| Data set | Sequencing specification | Sequencing NCBI accession | Sequencing depth<sup>a</sup> | Read length (bp)<sup>b</sup> | Number of reads | Reference genome | Genome length (Mbp) | Reference NCBI accession |
| --- | --- | --- | --- | --- | --- | --- | --- | --- |
| D1-I | Illumina Miseq | _<sup>c</sup> | 373x | 2×151 | 2×5 729 470 | E. coli K-12 MG1655 | 4.6 | NC_000913.3 |
| D1-P | Pacbio P6C4 | _<sup>d</sup> | 161x | 13 982 | 87 217 |  |  |  |
| D1-O | MinION R9 1D | _<sup>e</sup> | 319x | 14 891 | 164 472 |  |  |  |
| D2-I | Illumina Miseq | ERR1938683 | 81x | 2×150 | 2×3 318 467 | S. cerevisiae S288c | 12.2 | GCF_000146045.2 |
| D2-P | Pacbio P6C4 | PRJEB7245 | 120x | 8656 | 239 408 |  |  |  |
| D2-O | MinION R9 2D | ERP016443 | 59x | 7001 | 119 955 |  |  |  |
| D3-I | Illumina Nextseq | SRX3676782 | 44x | 2×151 | 2×20 619 401 | D. melanogaster ISO1 | 143.7 | GCF_000001215.4 |
| D3-P | Pacbio P5C3 | SRX499318 | 204x | 15 132 | 6 864 972 |  |  |  |
| D3-O | MinION R9.5 1D | SRX3676783 | 32x | 11 934 | 663 784 |  |  |  |

<sup>a</sup>Sequencing depth is estimated using the sequencing data and reference genome size
<sup>b</sup>N50 is reported for PacBio or ONT reads, since their lengths vary
<sup>c</sup>Downloaded from Illumina at [ftp://webdata.webdata@ussd-ftp.illumina.com/Data/SequencingRuns/MG1655/MiSeq\\_Ecoli\\_MG1655\\_110721\\_FF\\_R1.fastq.gz](ftp:///Data/SequencingRuns/MG1655/MiSeq_Ecoli_MG1655_110721_FF_R1.fastq.gz) and [ftp://webdata.webdata@ussd-ftp.illumina.com/Data/SequencingRuns/MG1655/MiSeq\\_Ecoli\\_MG1655\\_110721\\_FF\\_R2.fastq.gz](ftp:///Data/SequencingRuns/MG1655/MiSeq_Ecoli_MG1655_110721_FF_R2.fastq.gz)
<sup>d</sup>Downloaded from PacBio at https://github.com/PacificBiosciences/DevNet/wiki/E.-coli-Bacterial-Assembly
<sup>e</sup>Downloaded from Loman Labs at https://s3.climb.ac.uk/nanopore/E_coli_K12_1D_R9.2_SpotON_2.pass.fasta

To conduct performance evaluation under different sequencing depths, yeast sequencing reads (D2-P and D2-O) were subsampled randomly using Seqtk (https://github.com/lh3/seqtk). Subsamples with average depth of 10x, 20x and 30x were generated for ONT reads. In addition, 10x, 20x, 30x, 60x and 90x PacBio read subsamples were generated from D2-P. Details of these subsamples are available in Additional file 1 Table S1.

### Evaluation methodology

Our evaluation method takes corrected reads and a reference genome as input. The corrected reads were filtered using a user defined length (default 500). Reads which were too short to include in downstream analysis were dropped during the filtration. Filtered reads were aligned to the reference genome using Minimap2 [45] (using “-ax map-pb” and “-ax map-ont” parameters for PacBio and ONT reads respectively). The primary alignment for each read was used in the evaluation.

In an ideal scenario, an error correction software should take each erroneous long read and produce the error-free version of it, preserving each read and its full length. To assess how close to the ideal one can get, measures such as error rate post-correction or percentage of errors removed (termed *gain*; see [15]) can be utilized. However, long read error correction programs do not operate in this fashion. They may completely discard some reads or choose to split an input read into multiple reads when the high error rate cannot be reckoned with. In addition, short read assembly based error correction programs use long read alignments to de Bruijn graphs, and produce sequences corresponding to the aligned de Bruijn graph paths as output reads instead. Though original reads may not be fully preserved, all that matters for effective use of error correction software is that its output consists of sufficient number of high quality long reads that reflect adequate read lengths, sequencing depth, and coverage of the genome. Accordingly, our evaluation methodology reflects such assessment.

We measure the number of reads and total bases output by each error correction software, along with the number of aligned reads and total number of aligned bases extracted from alignment statistics, because they together reveal the effectiveness of correction. Besides, statistics which convey read length distribution such as maximum length and N50 were calculated to assess effect of the correction process on read lengths. Fraction of the genome covered by output reads is also reported to assess if there are regions of the genome that lost coverage or suffered significant deterioration in coverage depth post-correction. Any significant drop on these metrics can be a potential sign of information loss during the correction. Finally, alignment identity is calculated by the number of total matched bases divided by the total alignment length. Tools which achieve maximum alignment identity with minimum loss of information are desirable.

As part of this study, we provide an evaluation tool to automatically generate the evaluation statistics of corrected reads mentioned above (https://github.com/haowenz/LRECE). We include a wrapper script which can run state-of-the-art error correction software on a grid engine given any input data from user. Using the script, two types of evaluations can be conducted; users can either evaluate the performance on a list of tools with their own data to find a suitable tool for their studies, or they can run any correction tool with the benchmark data and compare it with other state-of-the-art tools.

## Results and discussion

### Experimental setup

All tests were run on the Swarm cluster located at Georgia Institute of Technology. Each compute node in the cluster has dual Intel Xeon CPU E5-2680 v4 (2.40 GHz) processors equipped with a total of 28 cores and 256GB main memory.

### Evaluated software

We evaluated 14 long read error correction programs in this study: Hercules, HG-CoLoR, FMLRC, HALC, CoLoRMap, Jabba, Nanocorr, proovread, LoRDEC, ECTools, LSC, FLAS, LoRMA and the error correction module in Canu. NaS was not included in the evaluation because it requires Newbler assembler which is no longer available from 454. PacBioToCA was also excluded since it is deprecated and no longer being maintained. The command line parameters were chosen based on user documentations of each software (Additional file 1 section “Versions and configurations”). The tools were configured to run exclusively on a single compute node and allowed to leverage all the 28 cores if multi-threading is supported. A cutoff on wall time was set to three days.

### Performance on benchmark data sets

We evaluated the quality and computational resource requirements of each software on our benchmark data sets. Results for the different data sets are shown in Tables 2, 3, 4, 5, 6 and 7. Because multiple factors are at play when considering accuracy, it is important to consider their collective influence in assessing quality of error correction. In what follows, we present a detailed and comparative discussion on correction accuracy, runtime and memory-usage. In addition, to guide error correction software users and future developers, we provide further insights into the strengths and limitations of various approaches that underpin the software. This includes evaluating their resilience to handle various sequencing depths, studying the effect of discarding or trimming input reads to gain higher accuracy, and impact on genome assembly.

**Table 2.**
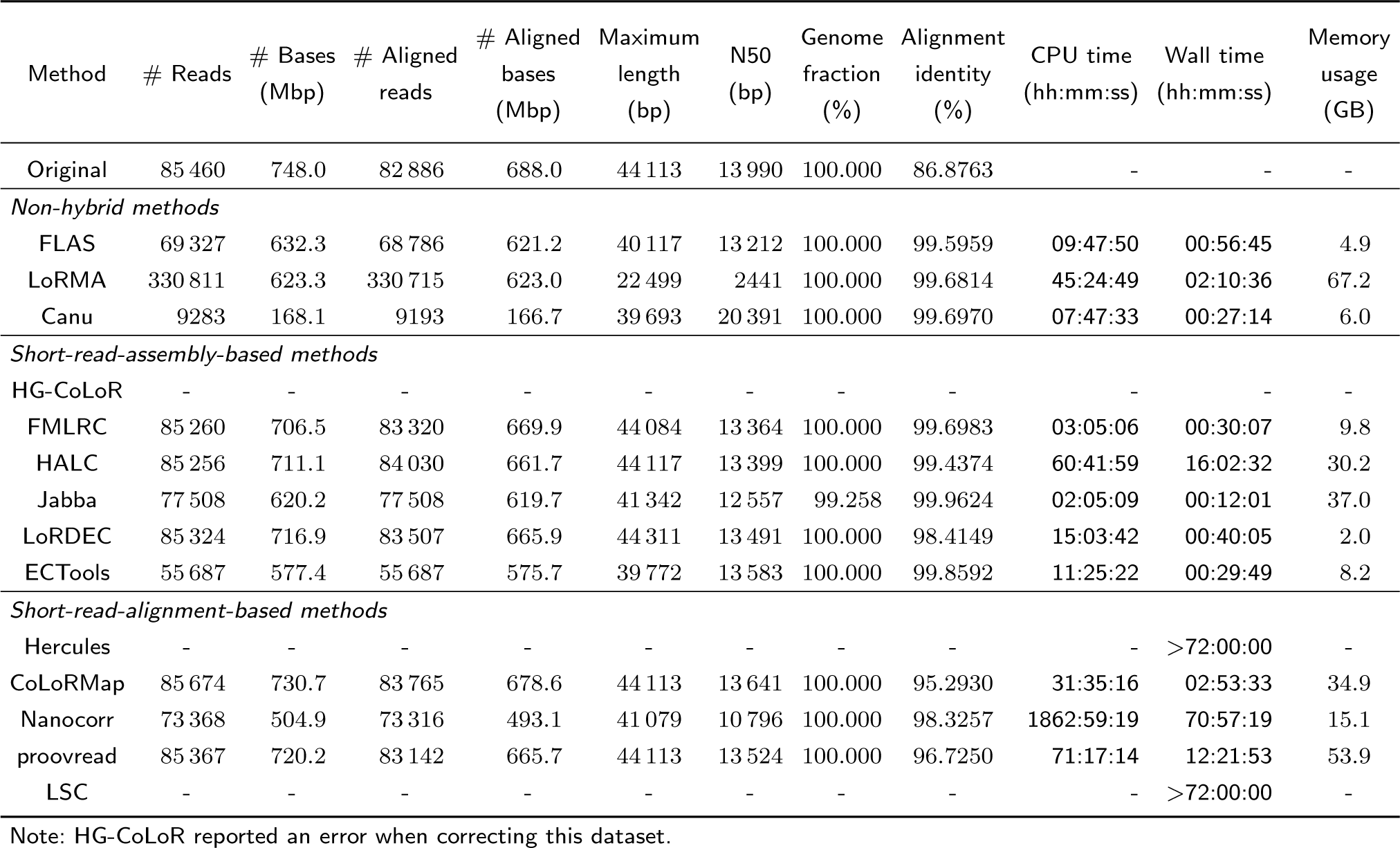
Experimental results for E. coli PacBio data set D1-P.

**Table 3.**
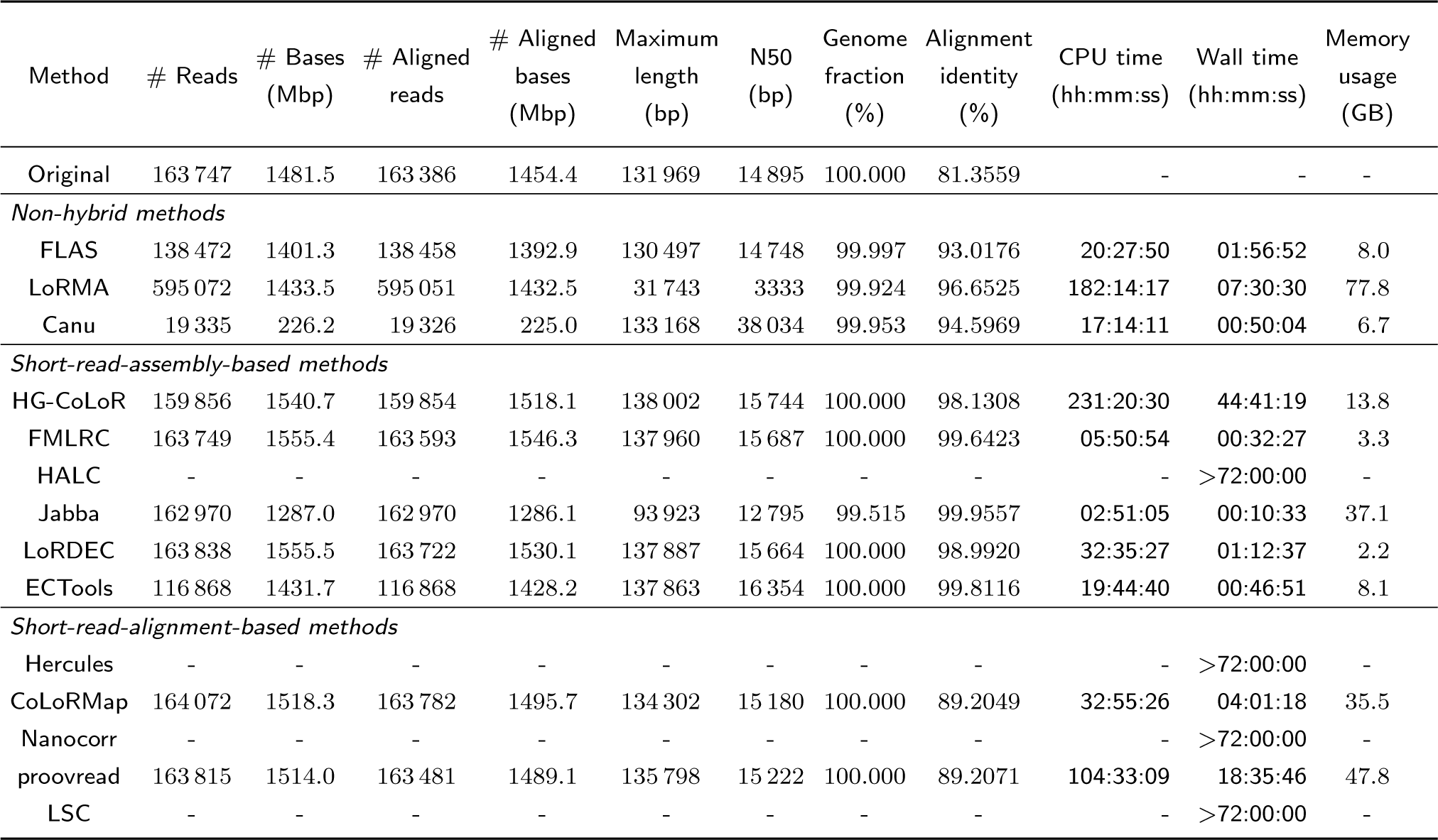
Experimental results for E. coli ONT data set D1-O.

**Table 4.**
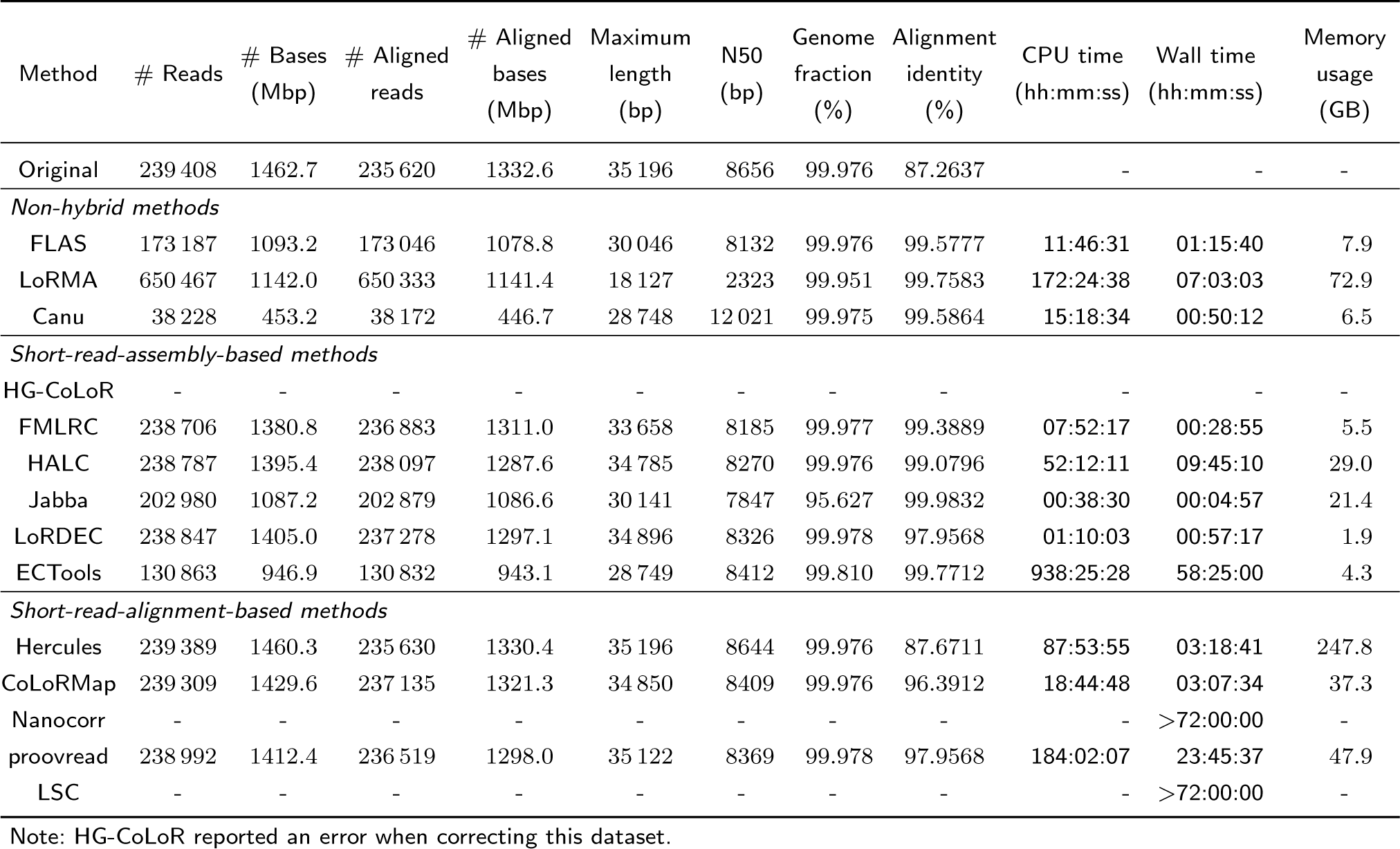
Experimental results for yeast PacBio data set D2-P.

**Table 5.**
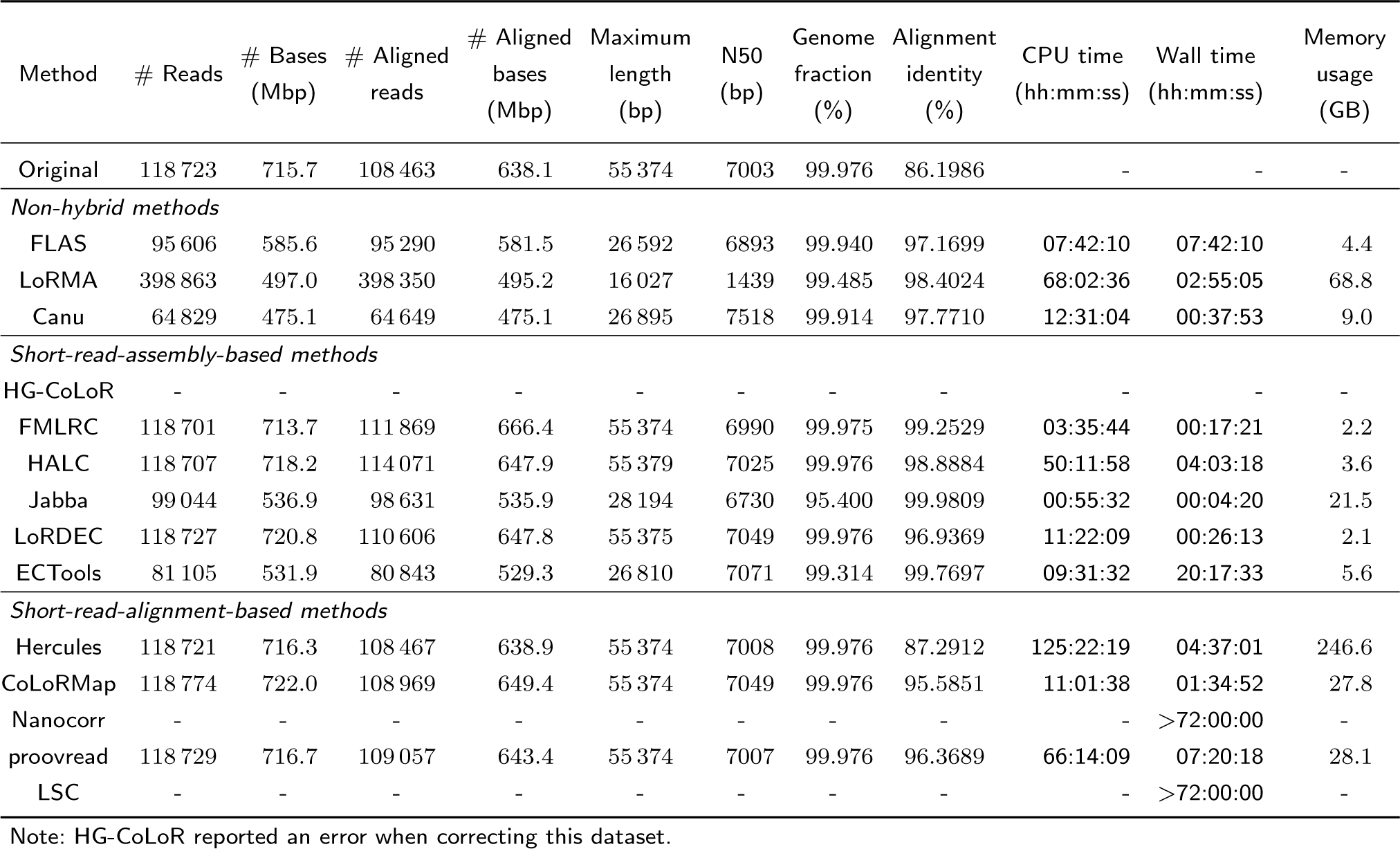
Experimental results for yeast ONT data set D2-O.

**Table 6.** Experimental results for fruit fly PacBio data set D3-P.

| Method | # Reads | # Bases (Mbp) | # Aligned reads | # Aligned bases (Mbp) | Maximum length (bp) | N50 (bp) | Genome fraction (%) | Alignment identity (%) | CPU time (hh:mm:ss) | Wall time (hh:mm:ss) | Memory usage (GB) |
| --- | --- | --- | --- | --- | --- | --- | --- | --- | --- | --- | --- |
| Original | 5 366 088 | 28 797.8 | 1 839 681 | 16 543.5 | 74 735 | 15 374 | 99.191 | 85.2734 | - | - | - |
| *Non-hybrid methods* |  |  |  |  |  |  |  |  |  |  |  |
| FLAS | 1 435 682 | 14 585.2 | 1 428 018 | 13 574.1 | 43 556 | 13 550 | 98.915 | 98.8363 | 271:44:27 | 36:30:42 | 53.1 |
| LoRMA | - | - | - | - | - | - | - | - | - | - | - |
| Canu | - | - | - | - | - | - | - | - | - | >72:00:00 | - |
| *Short-read-assembly-based methods* |  |  |  |  |  |  |  |  |  |  |  |
| HG-CoLoR | - | - | - | - | - | - | - | - | - | - | - |
| FMLRC | 5 246 485 | 27 354.6 | 2 477 890 | 16 543.5 | 74 735 | 14 554 | 99.191 | 96.5284 | 327:37:22 | 13:49:04 | 31.2 |
| HALC | 4 451 474 | 21 997.5 | 3 434 779 | 12 793.3 | 74 735 | 14 349 | 99.178 | 96.8863 | 770:35:46 | 55:58:24 | 73.0 |
| Jabba | 35 549 | 239.8 | 35 505 | 239.1 | 37 729 | 10 461 | 65.616 | 99.9615 | 656:05:15 | 24:33:41 | 175.8 |
| LoRDEC | 5 363 998 | 28 354.1 | 2 056 812 | 15 636.9 | 74 719 | 15 078 | 99.200 | 92.2954 | 1011:52:27 | 36:19:18 | 5.9 |
| ECTools | - | - | - | - | - | - | - | - | - | >72:00:00 | - |
| *Short-read-alignment-based methods* |  |  |  |  |  |  |  |  |  |  |  |
| Hercules | - | - | - | - | - | - | - | - | - | - | - |
| CoLoRMap | 5 366 107 | 28 891.6 | 1 841 822 | 14 976.8 | 74 735 | 15 442 | 99.189 | 83.2580 | 495:11:17 | 64:52:25 | 189.4 |
| Nanocorr | - | - | - | - | - | - | - | - | - | >72:00:00 | - |
| proovread | - | - | - | - | - | - | - | - | - | >72:00:00 | - |
| LSC | - | - | - | - | - | - | - | - | - | >72:00:00 | - |

Note: LoRMA, HG-CoLoR and Hercules reported errors when correcting this dataset.

**Table 7.** Experimental results for fruit fly ONT data set D3-O.

| Method | # Reads | # Bases (Mbp) | # Aligned reads | # Aligned bases (Mbp) | Maximum length (bp) | N50 (bp) | Genome fraction (%) | Alignment identity (%) | CPU time (hh:mm:ss) | Wall time (hh:mm:ss) | Memory usage (GB) |
| --- | --- | --- | --- | --- | --- | --- | --- | --- | --- | --- | --- |
| Original | 642 255 | 4609.5 | 554 083 | 3857.9 | 446 050 | 11 956 | 98.719 | 83.5921 | - | - | - |
| *Non-hybrid methods* |  |  |  |  |  |  |  |  |  |  |  |
| FLAS | 423 097 | 3507.6 | 422 206 | 3402.6 | 64 365 | 11 517 | 97.588 | 95.3301 | 23:04:50 | 03:12:50 | 10.8 |
| LoRMA | 703 097 | 615.5 | 682 288 | 592.3 | 32 644 | 865 | 30.338 | 98.1230 | 666:37:35 | 25:52:14 | 92.8 |
| Canu | 430 082 | 3415.6 | 421 475 | 3220.2 | 254 967 | 12 090 | 97.592 | 96.3739 | 88:51:10 | 04:36:20 | 20.2 |
| *Short-read-assembly-based methods* |  |  |  |  |  |  |  |  |  |  |  |
| HG-CoLoR | - | - | - | - | - | - | - | - | - | - | - |
| FMLRC | 641 945 | 4647.2 | 578 290 | 3978.2 | 444 605 | 12 088 | 98.592 | 97.6010 | 47:45:17 | 03:06:05 | 31.2 |
| HALC | 643 002 | 4668.5 | 611 191 | 3955.7 | 451 284 | 12 115 | 98.616 | 97.6634 | 126:30:01 | 05:43:37 | 42.4 |
| Jabba | 494 546 | 2878.2 | 494 430 | 2876.3 | 72 501 | 9305 | 83.166 | 99.9745 | 175:19:34 | 06:56:29 | 136.8 |
| LoRDEC | 642 882 | 4655.9 | 567 878 | 3921.1 | 447 726 | 12 079 | 98.691 | 94.0382 | 152:05:32 | 05:38:05 | 5.7 |
| ECTools | - | - | - | - | - | - | - | - | - | >72:00:00 | - |
| *Short-read-alignment-based methods* |  |  |  |  |  |  |  |  |  |  |  |
| Hercules | 642 287 | 4612.8 | 554 630 | 3859.4 | 449 799 | 11 966 | 98.713 | 83.9340 | 398:10:17 | 17:32:36 | 247.7 |
| CoLoRMap | 649 041 | 4692.1 | 565 881 | 3963.8 | 442 948 | 12 050 | 98.715 | 94.3361 | 160:00:22 | 16:07:18 | 57.3 |
| Nanocorr | - | - | - | - | - | - | - | - | - | >72:00:00 | - |
| proovread | - | - | - | - | - | - | - | - | - | >72:00:00 | - |
| LSC | - | - | - | - | - | - | - | - | - | >72:00:00 | - |

Note: HG-CoLoR reported an error when correcting this dataset.

#### Correction quality

We measure quality using the number of reads and total bases output in comparison with the input, the resulting alignment identity, fraction of the genome covered and read length distribution including maximum size and N50 length. From Tables 2, 3, 4, 5, 6 and 7, we gather that the best performing hybrid methods (e.g., FMLRC) are capable of correcting reads to achieve base-level accuracy in the high 90’s. For the *E. coli* and yeast data sets, many of these programs achieve alignment identity > 99%. A crucial aspect to consider here is whether the high accuracy is achieved while preserving input read depth and N50. Few tools (e.g. Jabba) seem to attain high alignment identity at the cost of producing shorter reads and reduced depths because they choose to either discard uncorrected reads or trim the uncorrected regions. This may have a negative impact on downstream analyses. This trade-off is further discussed later in “Effect of discarding reads during correction” section.

Among the hybrid methods, a key observation is that short-read-assembly-based methods tend to show better performance than short-read-alignment-based methods. We provide the following explanation. Given that long reads are error-prone, short read alignment to long reads is more likely to be wrong (or ambiguous) than long read alignment to graph structures built using short reads. Errors in long reads can cause false positives in identifying the true positions where the respective short reads should align, which causes false correction later. For example, during the correction of D3-P, the alignment identity of corrected reads generated by CoLoRMap in fact decreased when compared to the uncorrected reads. The reason is that CoLoRMap uses BWA-mem [46] to map short reads, which is designed to report best mapping. However, due to the high error rates, the best mapping is not necessarily the true mapping. Large volume of erroneous long reads in D3-P can lead to many false alignments, which affected the correction process. On the other hand, long read lengths make it possible to have higher confidence when aligning them to paths in the graph. Therefore, in most of the experiments, assembly-based methods were able to produce reasonable correction. Non-hybrid correction is more challenging as it relies solely on overlaps between erroneous long reads, yet the tools in this category yield competitive accuracy in many cases. However, non-hybrid methods may achieve lower alignment identity when the long reads are more erroneous. For example, the alignment identity of corrected reads generated by FLAS, LoRMA and Canu is lower than almost all hybrid methods for D1-O where the average alignment identity of uncorrected reads is only 81.36%.

#### Runtime and memory usage

Scalability of the correction tools is an important aspect to consider in their evaluation. Slow speed or high memory usage makes it difficult to apply them to correct large data sets. Our results show that hybrid methods, in particular assembly-based methods, are much faster than the rest. For instance, Hercules and LSC failed to generate corrected reads in three days for D1-P, while most of the assembly-based tools finished the same job in less than one hour. Hercules, Nanocorr and LSC were unable to finish the correction of D2-O in three days, which was finished by FMLRC or LoRDEC in hours. Although proovread can complete the corrections of D2-P and D2-O, the run-time was 49.3 and 34.4 times longer, respectively, than runtime needed by FMLRC. Moreover, assembly-based methods, e.g., LoRDEC and FMLRC, used less memory in most of the experiments. Therefore, in terms of computational performance, users should give priority to short-read assembly-based methods over short-read alignment-based methods.

Among the non-hybrid methods, LoRMA’s memory usage was generally the highest among all the tools, and was slower than assembly-based methods. However, Canu showed superior scalability. Owing to a fast long read overlap detection algorithm using Min-Hash [44], Canu was able to compute long read overlaps and used them to correct the reads in reasonable time, which is comparable to most of the hybrid methods. The memory footprint of Canu was also lower than many hybrid-methods. However, Canu did not finish the correction of D3-P in three days probably because this data set is too large to compute pairwise overlaps. FLAS showed performance comparable to Canu as FLAS also leverages the fast overlap computation method in MECAT [43].

#### Effect of long read sequencing depth on error correction

Requiring high sequencing coverage for effective error correction can impact both cost and time consumed during sequencing and analysis. The relative cost per base pair using third-generation sequencing is still several folds higher when compared to the latest Illumina sequencers [1]. Accordingly, we study how varying long read sequencing depth affects correction quality, while keeping the short read data set fixed. We conducted this experiment using data sets D2-P and D2-O with various depth levels obtained using random sub-sampling. The details of the subsamples are summarized in Additional file 1 Table S1. The output behavior of the correction tools is shown in Additional file 1 Tables S2-S9.

For corrected reads generated by hybrid methods, no significant change on the metrics was observed except those generated by CoLoRMap. The alignment identity of its corrected reads increased with decreased sequencing depth. This observation is consistent with the experimental results reported by its authors. Similarly, CoLoRMap did not perform well on large data sets such as D3-P as large data sets increase the risk of false positive alignments.

On the other hand, the performance of non-hybrid methods deteriorated significantly when sequencing depth was decreased. As non-hybrid methods leverage overlap information to correct errors, they require sufficient long read coverage to make true correction. The genome fraction covered by corrected reads produced by LoRMA with subsamples of D2-P decreased from 99.59% to 82.97% when sequencing depth dropped from 90x to 60x, and further decreased to 9.61%, 5.39% and 3.78% for 30x, 20x and 10x respectively, implying loss of many long reads after correction. The alignment identities were still greater than 99% using all sub-samples because LoRMA trimmed the uncorrected regions. For corrected reads generate by Canu, no significant change on genome fraction was observed. But the alignment identity dropped from above 99% to 97.03% and 95.63% for 20x and 10x sequencing depths, respectively. FLAS showed similar performance but genome fraction for 10x sequencing depth was only 90.20% lower than the 99.92% achieved by Canu, which indicates FLAS drops some reads when sequencing depth is low.

#### Effect of discarding reads during correction

Many correction tools opt for discarding input reads or regions within reads that they fail to correct. As a result, the reported alignment identity is high (> 99%), but much fewer number of bases survive after correction. This effect is more pronounced in corrected reads generated by Jabba, ECTools, and LoRMA. They either trim uncorrected regions at sequence ends, or even in the middle, to avoid errors in the final output which eventually yields high alignment identity. However, aggressive trimming also makes the correction lossy and may influence downstream analysis because long range information is lost if the reads are shortened or broken into smaller pieces. Therefore, users should be conservative in trimming and turn it off when necessary. One good practice is to keep the uncorrected regions and let downstream analysis tools perform the trimming, e.g. overlap-based trimming after read correction in Canu. A direct implication of discarding or trimming reads is the change of read length distribution. Figure 1 shows the original and corrected read length distributions. Among all the tools, HG-CoLoR, FMLRC, HALC, CoLoRMAP, LoRDEC and proovread can maintain a similar read length distribution after correction whereas Nanocorr, Jabba and ECTools lost many long reads after correction due to their trimming step. Nanocorr drops a long read when there is no short read aligning to it. This procedure can remove many error-prone long reads, which leads to a higher alignment identity after correction. However, the fraction of discarded reads in many cases is found to be significant. For example, a mere 376.3 million bp cumulative length of sequences survived out of 1462.7 million bp data set, after correction of D2-P. ECTools also generated only 946.9 million corrected bases using this data set. Canu changed the read length distribution significantly after correction although due to a different reason. Canu estimates the read length after correction and tries to keep the longest 40x reads for subsequent assembly. FLAS kept most of the reads with short length while losing many reads with long length.

**Figure 1 E.**
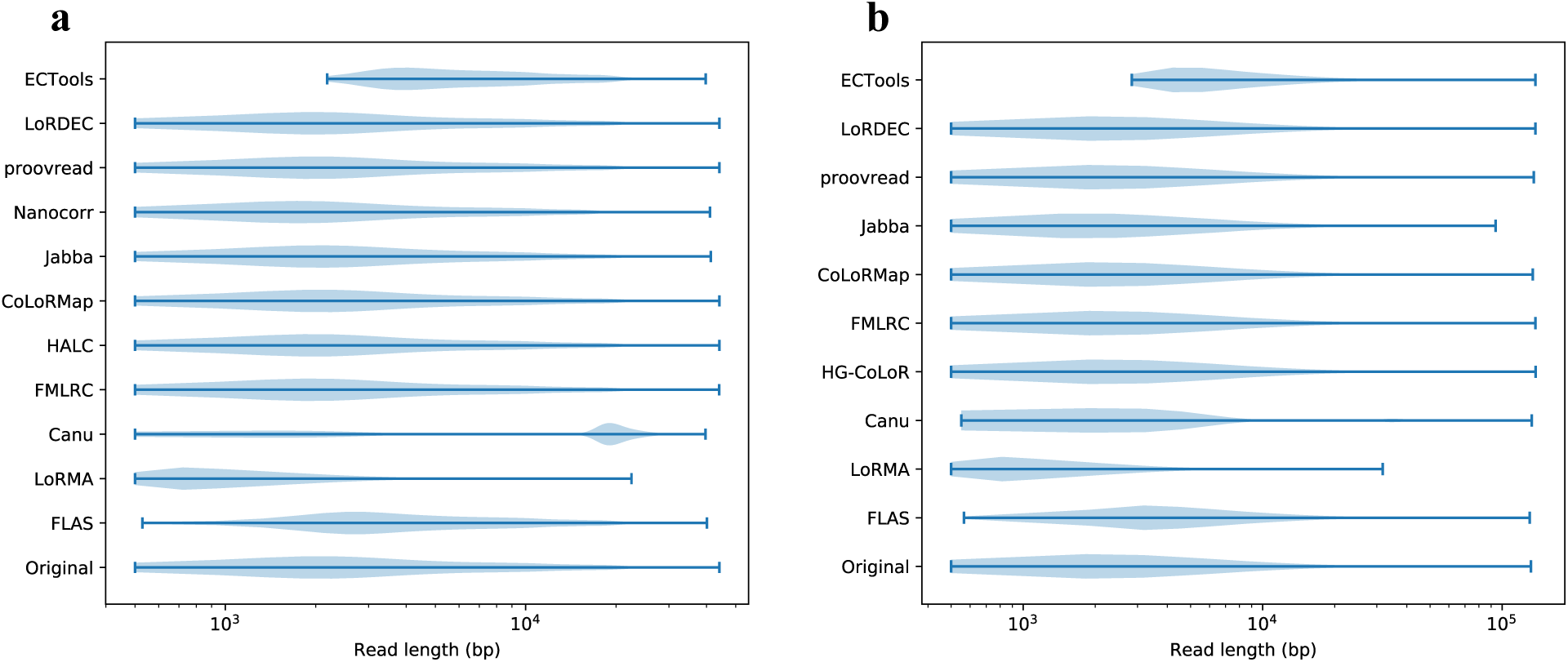
coli corrected read length distribution. Corrected read length distribution is shown in violinplots for *E. coli* PacBio and ONT sequencing data in **a** and **b** respectively. Note that ECTools only corrects reads longer than 1000 bp and drops the reads shorter than that.

#### Effect of error correction on genome assembly

Error correction of long reads remains a useful pre-processing stage for reliable construction of overlap graphs during genome assembly. We examined how well the accuracy of error correction correlates with the quality of genome assembly performed using corrected reads. To do so, we conducted an experiment to compute genome assembly using corrected PacBio and ONT reads of *E. coli*, i.e., corrected reads for D1-P and D1-O. Assembly was computed using Canu with its error correction module turned off, and assembly quality was assessed using QUAST [47]; the results are shown in Tables 8 and 9.

**Table 8.** Results of genome assembly computed using corrected reads of D1-P.

| Method | # contigs | NGA50 (bp) | Largest contigs (bp) | Total length (bp) | Genome fraction (%) | # misassemblies | # mismatches | # indels (<=5bp) | # indels (>5bp) | Indel length (bp) |
| --- | --- | --- | --- | --- | --- | --- | --- | --- | --- | --- |
| *Non-hybrid methods* |  |  |  |  |  |  |  |  |  |  |
| FLAS | 2 | 3 996 362 | 4 681 650 | 4 689 583 | 99.998 | 4 | 4 | 162 | 0 | 167 |
| LoRMA | 14 | 696 878 | 2 501 146 | 4 663 900 | 99.938 | 4 | 75 | 4181 | 6 | 4295 |
| Canu | 1 | 3 976 437 | 4 670 120 | 4 670 120 | 99.998 | 4 | 7 | 92 | 0 | 95 |
| *Short-read-assembly-based methods* |  |  |  |  |  |  |  |  |  |  |
| FMLRC | 9 | 3 821 409 | 4 657 352 | 4 831 908 | 99.998 | 8 | 1 | 4 | 0 | 5 |
| HALC | 25 | 2 947 777 | 4 682 714 | 5 388 722 | 99.983 | 8 | 541 | 35 | 8 | 257 |
| Jabba | 58 | 138 874 | 398 327 | 4 623 296 | 97.273 | 1 | 172 | 32 | 3 | 167 |
| LoRDEC | 2 | 3 996 441 | 4 681 757 | 4 703 690 | 99.998 | 4 | 66 | 18 | 2 | 55 |
| ECTools | 19 | 3 548 731 | 4 657 296 | 5 154 324 | 99.974 | 4 | 592 | 80 | 2 | 188 |
| *Short-read-alignment-based methods* |  |  |  |  |  |  |  |  |  |  |
| CoLoRMap | 86 | 1 217 587 | 1 448 649 | 5 700 143 | 99.998 | 4 | 42 | 3 | 7 | 478 |
| Nanocorr | 18 | 3 095 077 | 4 646 253 | 4 931 697 | 99.998 | 5 | 65 | 34 | 2 | 157 |
| proovread | 2 | 1 686 030 | 4 626 702 | 4 666 724 | 99.656 | 4 | 76 | 97 | 1 | 176 |

**Table 9.** Results of genome assembly computed using corrected reads of D1-O.

| Method | # contigs | NGA50 (bp) | Largest contigs (bp) | Total length (bp) | Genome fraction (%) | # misassemblies | # mismatches | # indels (<=5bp) | # indels (>5bp) | Indel length (bp) |
| --- | --- | --- | --- | --- | --- | --- | --- | --- | --- | --- |
| *Non-hybrid methods* |  |  |  |  |  |  |  |  |  |  |
| FLAS | 1 | 1 283 465 | 4 561 925 | 4 561 925 | 99.834 | 4 | 4785 | 70 062 | 1310 | 122 264 |
| LoRMA | 14 | 726 649 | 1 239 048 | 4 583 602 | 99.650 | 3 | 23 275 | 57 650 | 37 | 73 578 |
| Canu | 1 | 3 335 496 | 4 601 279 | 4 601 279 | 99.914 | 2 | 14 810 | 56 511 | 283 | 78 931 |
| *Short-read-assembly-based methods* |  |  |  |  |  |  |  |  |  |  |
| HG-CoLoR | 32 | 3 924 167 | 4 634 988 | 5 555 776 | 99.845 | 28 | 393 | 82 | 8 | 246 |
| FMLRC | 9 | 4 325 756 | 4 718 452 | 4 974 808 | 99.874 | 6 | 6 | 7 | 4 | 242 |
| Jabba | 57 | 105 474 | 311 624 | 4 460 218 | 95.838 | 0 | 117 | 22 | 5 | 179 |
| LoRDEC | 57 | 3 492 326 | 4 044 623 | 5 389 657 | 99.800 | 10 | 402 | 162 | 18 | 448 |
| ECTools | 2 | 2 891 718 | 4 733 248 | 4 797 686 | 99.885 | 3 | 632 | 267 | 16 | 682 |
| *Short-read-alignment-based methods* |  |  |  |  |  |  |  |  |  |  |
| CoLoRMap | 55 | 206 971 | 501 963 | 6 007 440 | 94.790 | 4 | 11 074 | 14 392 | 206 | 24 183 |
| proovread | 99 | 75 162 | 225 568 | 4 613 429 | 98.737 | 14 | 276 | 41 | 1 | 75 |

Considering the assemblies generated using corrected PacBio reads (Table 8), NGA50 of about 3 Mbp was obtained when using reads generated by FLAS, Canu, FMLRC, HALC, Nanocorr, LoRDEC or ECTools. When using corrected ONT reads (Table 9), assemblies generated using reads corrected by Canu, HG-CoLoR, FMLRC, LoRDEC and ECTools have NGA50 near 3 Mbp. In contrast, assemblies generated using reads corrected by Jabba and LoRMA showed lower NGA50 in both cases. Their trimming procedure possibly led to the loss of some long range information, thereby causing lower continuity in assembly. Post error correction, alignment identity of corrected reads needs to be sufficiently high to identify true overlaps during assembly. We observe that NGA50 of assemblies generated using reads corrected by CoLoRMap and proovread is low, as the corrected reads generated by these two tools have low alignment identity (e.g.,< 90% for D1-O, Table 3).

We also examined the frequency of mismatches and indels in the assemblies. For data set D1-P, corrected reads generated by HALC and ECTools produced assemblies with > 500 mismatches, significantly higher than the other tools. However, alignment identity of their corrected reads was either competitive with, or superior to, what is produced by other tools. Notably, both HALC and ECTools use assembled contigs from short reads to do error correction. Mis-assemblies of short reads, especially in repetitive and low-complexity regions, may cause false corrections, which leads to errors during assembly [29]. Corrected reads produced by FMLRC achieved the least number of errors in assembly. Meanwhile, its alignment identity was also the highest among the methods which avoid trimming. Therefore, higher alignment identity of corrected reads is an important but not a sufficient criteria to minimize errors in genome assemblies.

Non-hybrid methods such as LoRMA, Canu and FLAS produced more indels than mismatches in their assemblies while most of the hybrid methods showed the opposite behavior. These observations suggest that existing self-correction methods are not good at handling indels when compared to hybrid methods. Consequently, *de novo* long read assemblers that use self-correction methods typically rely on post-processing ‘polishing’ stages, using signal-level data from long read instruments [2, 33] or alternate sequencing technologies [48].

## Conclusions and future directions

In this work, we established benchmark data sets and evaluation methods for comprehensive assessment of long read error correction software. Our results suggest that hybrid methods aided by short accurate reads can achieve better correction quality, especially when handling low coverage-depth long reads,compared with non-hybrid methods. Within the hybrid methods, assembly-based methods are superior to alignment-based methods in terms of scalability to large data sets. Besides, better performance on correction such as preserving higher proportion of input bases and high alignment identity often leads to better performance in downstream applications such as genome assembly. But the tools with superior correction performance should be further tested in the context of applications of interest.

Users can also select tools according to our experimental results for their specific expectations. When speed is a concern, assembly-based hybrid methods are preferred whenever short reads are available. Besides, hybrid methods are more immune to low long read sequencing depth than non-hybrid methods. Thus, users are recommended to choose hybrid methods when long read sequencing depth is relatively low. In cases where indel errors may cause a serious negative impact on downstream analyses, hybrid methods should be preferred over non-hybrid ones if short reads are available.

FMLRC outperformed other hybrid methods in almost all the experiments. For non-hybrid methods, Canu and FLAS showed better performance over LoRMA. Hence, these three are recommended as default when users want to avoid laborious tests on all the error correction tools.

For future work, better self-correction algorithms are expected to avoid hybrid sequencing, thus reducing experimental labor on short read sequencing preparation. In addition, most of the correction algorithms run for days to correct errors in the sequencing of even moderately large and complex genomes like the fruit fly. These algorithms will spend much more time on correcting larger sequencing data sets of human, and become a bottleneck in sequencing data analysis. Therefore, more efficient or parallel correction algorithms should be developed to ease the computational burden. Furthermore, none of the hybrid tools makes use of paired-end information in their correction, except CoLoRMap. But the use of paired-end reads in CoLoRMap did not improve correction performance significantly according to previous studies. Paired-end reads have already been used to resolve repeats and remove entanglements in de Bruijn graphs [49]. Since many error correction tools build de Bruijn graphs to correct long reads, the paired-end information may also be able to improve error correction.

Most of the published error correction tools focus on correction of long DNA reads sequenced from a single genome, which also served as the motivation for our review. Long read sequencing is increasingly gaining traction in transcriptomics and metagenomics applications. It is not clear whether the existing tools can be leveraged or extended to work effectively in such scenarios, and is an active area of research [50].

## Declarations

### Ethics approval and consent to participate

Not applicable.

### Consent for publication

Not applicable.

### Availability of data and materials

All data generated or analysed during this study are included in this published article (and its supplementary information files).

### Author’s contributions

HZ, CJ and SA designed the study. HZ performed the experiments and wrote the paper. CJ and SA revised the paper. All authors read and approved the final manuscript.

## Acknowledgements

Not applicable.

## Additional Files

### Additional file 1

Settings of the error correction tools and correction results on subsamples of yeast PacBio and ONT sequencing data.

## References

1. Sedlazeck FJ, Lee H, Darby CA, Schatz MC. Piercing the dark matter: bioinformatics of long-range sequencing and mapping. Nature Reviews Genetics. 2018;p. 1.

2. Loman NJ, Quick J, Simpson JT. A complete bacterial genome assembled de novo using only nanopore sequencing data. Nature methods. 2015;12(8):733.

3. Chin CS, Peluso P, Sedlazeck FJ, Nattestad M, Concepcion GT, Clum A, et al. Phased diploid genome assembly with single-molecule real-time sequencing. Nature methods. 2016;13(12):1050.

4. Jain M, Koren S, Miga KH, Quick J, Rand AC, Sasani TA, et al. Nanopore sequencing and assembly of a human genome with ultra-long reads. Nature biotechnology. 2018;36(4):338.

5. Sedlazeck FJ, Rescheneder P, Smolka M, Fang H, Nattestad M, von Haeseler A, et al. Accurate detection of complex structural variations using single molecule sequencing. Preprint at https://www.biorxivorg/content/arly/2017/07/28/169557. 2017;.

6. Chaisson MJ, Huddleston J, Dennis MY, Sudmant PH, Malig M, Hormozdiari F, et al. Resolving the complexity of the human genome using single-molecule sequencing. Nature. 2015;517(7536):608.

7. Gordon SP, Tseng E, Salamov A, Zhang J, Meng X, Zhao Z, et al. Widespread polycistronic transcripts in fungi revealed by single-molecule mRNA sequencing. PloS one. 2015;10(7):e0132628.

8. Dilthey A, Jain C, Koren S, Phillippy A. MetaMaps-Strain-level metagenomic assignment and compositional estimation for long reads. bioRxiv. 2018;p. 372474.

9. Rand AC, Jain M, Eizenga JM, Musselman-Brown A, Olsen HE, Akeson M, et al. Mapping DNA methylation with high-throughput nanopore sequencing. Nature methods. 2017;14(4):411.

10. Simpson JT, Workman RE, Zuzarte P, David M, Dursi L, Timp W. Detecting DNA cytosine methylation using nanopore sequencing. Nature methods. 2017;14(4):407.

11. Carneiro MO, Russ C, Ross MG, Gabriel SB, Nusbaum C, DePristo MA. Pacific biosciences sequencing technology for genotyping and variation discovery in human data. BMC genomics. 2012;13(1):375.

12. Jain M, Fiddes IT, Miga KH, Olsen HE, Paten B, Akeson M. Improved data analysis for the MinION nanopore sequencer. Nature methods. 2015;12(4):351.

13. Korlach J, Biosciences P. Understanding Accuracy in SMRT® Sequencing; 2013.

14. Ashton PM, Nair S, Dallman T, Rubino S, Rabsch W, Mwaigwisya S, et al. MinION nanopore sequencing identifies the position and structure of a bacterial antibiotic resistance island. Nature biotechnology. 2015;33(3):296.

15. Yang X, Chockalingam SP, Aluru S. A survey of error-correction methods for next-generation sequencing. Briefings in bioinformatics. 2012;14(1):56–66.

16. Alic AS, Ruzafa D, Dopazo J, Blanquer I. Objective review of de novo stand-alone error correction methods for NGS data. Wiley Interdisciplinary Reviews: Computational Molecular Science. 2016;6(2):111–146.

17. Koren S, Schatz MC, Walenz BP, Martin J, Howard JT, Ganapathy G, et al. Hybrid error correction and de novo assembly of single-molecule sequencing reads. Nature biotechnology. 2012;30(7):693.

18. Au KF, Underwood JG, Lee L, Wong WH. Improving PacBio long read accuracy by short read alignment. PloS one. 2012;7(10):e46679.

19. Lee H, Gurtowski J, Yoo S, Marcus S, McCombie WR, Schatz M. Error correction and assembly complexity of single molecule sequencing reads. BioRxiv. 2014;p. 006395.

20. Salmela L, Rivals E. LoRDEC: accurate and efficient long read error correction. Bioinformatics. 2014;30(24):3506–3514.

21. Hackl T, Hedrich R, Schultz J, Förster F. proovread: large-scale high-accuracy PacBio correction through iterative short read consensus. Bioinformatics. 2014;30(21):3004–3011.

22. Madoui MA, Engelen S, Cruaud C, Belser C, Bertrand L, Alberti A, et al. Genome assembly using Nanopore-guided long and error-free DNA reads. BMC genomics. 2015;16(1):327.

23. Goodwin S, Gurtowski J, Ethe-Sayers S, Deshpande P, Schatz MC, McCombie WR. Oxford Nanopore sequencing, hybrid error correction, and de novo assembly of a eukaryotic genome. Genome research. 2015;25(11):1750–1756.

24. Miclotte G, Heydari M, Demeester P, Rombauts S, Van de Peer Y, Audenaert P, et al. Jabba: hybrid error correction for long sequencing reads. Algorithms for Molecular Biology. 2016;11(1):10.

25. Haghshenas E, Hach F, Sahinalp SC, Chauve C. Colormap: Correcting long reads by mapping short reads. Bioinformatics. 2016;32(17):i545–i551.

26. Salmela L, Walve R, Rivals E, Ukkonen E. Accurate self-correction of errors in long reads using de Bruijn graphs. Bioinformatics. 2016;33(6):799–806.

27. Bao E, Lan L. HALC: High throughput algorithm for long read error correction. BMC bioinformatics. 2017;18(1):204.

28. Bao E, Xie F, Song C, Dandan S. Hals: Fast and high throughput algorithm for pacbio long read self-correction. RECOMB-SEQ; 2018.

29. Wang JR, Holt J, McMillan L, Jones CD. FMLRC: Hybrid long read error correction using an FM-index. BMC bioinformatics. 2018;19(1):50.

30. Morisse P, Lecroq T, Lefebvre A, Berger B. Hybrid correction of highly noisy long reads using a variable-order de Bruijn graph. Bioinformatics. 2018;.

31. Firtina C, Bar-Joseph Z, Alkan C, Cicek AE. Hercules: a profile HMM-based hybrid error correction algorithm for long reads. Nucleic acids research. 2018;46(21):e125–e125.

32. Koren S, Walenz BP, Berlin K, Miller JR, Bergman NH, Phillippy AM. Canu: scalable and accurate long-read assembly via adaptive k-mer weighting and repeat separation. Genome research. 2017;27(5):722–736.

33. Chin CS, Alexander DH, Marks P, Klammer AA, Drake J, Heiner C, et al. Nonhybrid, finished microbial genome assemblies from long-read SMRT sequencing data. Nature methods. 2013;10(6):563.

34. Laehnemann D, Borkhardt A, McHardy AC. Denoising DNA deep sequencing data high-throughput sequencing errors and their correction. Briefings in bioinformatics. 2015;17(1):154–179.

35. Mahmoud M, Zywicki M, Twardowski T, Karlowski WM. Efficiency of PacBio long read correction by 2nd generation Illumina sequencing. Genomics. 2017;.

36. La S, Haghshenas E, Chauve C. LRCstats, a tool for evaluating long reads correction methods. Bioinformatics. 2017;33(22):3652–3654.

37. Fichot EB, Norman RS. Microbial phylogenetic profiling with the Pacific Biosciences sequencing platform. Microbiome. 2013;1(1):10.

38. Stöcker BK, Köster J, Rahmann S. SimLoRD: simulation of long read data. Bioinformatics. 2016;32(17):2704–2706.

39. Altschul SF, Gish W, Miller W, Myers EW, Lipman DJ. Basic local alignment search tool. Journal of molecular biology. 1990;215(3):403–410.

40. Pop M, Phillippy A, Delcher AL, Salzberg SL. Comparative genome assembly. Briefings in bioinformatics. 2004;5(3):237–248.

41. Yang X, Dorman KS, Aluru S. Reptile: representative tiling for short read error correction. Bioinformatics. 2010;26(20):2526–2533.

42. Kowalski T, Grabowski S, Deorowicz S. Indexing arbitrary-length k-mers in sequencing reads. PloS one. 2015;10(7):e0133198.

43. Xiao CL, Chen Y, Xie SQ, Chen KN, Wang Y, Han Y, et al. MECAT: fast mapping, error correction, and de novo assembly for single-molecule sequencing reads. nature methods. 2017;14(11):1072.

44. Berlin K, Koren S, Chin CS, Drake JP, Landolin JM, Phillippy AM. Assembling large genomes with single-molecule sequencing and locality-sensitive hashing. Nature biotechnology. 2015;33(6):623.

45. Li H. Minimap2: pairwise alignment for nucleotide sequences. Bioinformatics. 2018;1:7.

46. Li H. Aligning sequence reads, clone sequences and assembly contigs with BWA-MEM. arXiv preprint arXiv:13033997. 2013;.

47. Gurevich A, Saveliev V, Vyahhi N, Tesler G. QUAST: quality assessment tool for genome assemblies. Bioinformatics. 2013;29(8):1072–1075.

48. Walker BJ, Abeel T, Shea T, Priest M, Abouelliel A, Sakthikumar S, et al. Pilon: an integrated tool for comprehensive microbial variant detection and genome assembly improvement. PloS one. 2014;9(11):e112963.

49. Bankevich A, Nurk S, Antipov D, Gurevich AA, Dvorkin M, Kulikov AS, et al. SPAdes: a new genome assembly algorithm and its applications to single-cell sequencing. Journal of computational biology. 2012;19(5):455–477.

50. de Lima LIS, Marchet C, Caboche S, Da Silva C, Istace B, Aury JM, et al. Comparative assessment of long-read error-correction software applied to RNA-sequencing data. bioRxiv. 2018;p. 476622.

