## Supplementary for "A comprehensive evaluation of long read error correction methods"

### A comprehensive evaluation of long read error correction methods: Supplementary material

### Versions and configurations

The version and configuration for each error correction tool are as follows.

- Hercules: version 0.1 was used. Bowtie2 [1] was used to align compressed short reads to compressed long reads, which is suggested in the manual. The default parameters were used.
- HG-CoLoR: commit cc18a95 was used. The maximum K-mer size (–maxK) of the variable-order de Bruijn graph was set to 100 as suggested in its paper. We set top alignments reported by BLASR [2] (–bestn) to 30, 40 and 50 for *E. coli*, yeast and fruit fly data sets respectively, according to the short reads sequencing depth, which is recommended in its manual.
- FMLRC: version 0.1.2 was used. Before running FMLRC, a BWT of short reads was constructed with msBWT [3] version 0.3.0 using default parameters according to its manual. FMLRC was run with default short and long k-mer length, which is 21 and 59 respectively.
- HALC: version 1.1 was used. Prior to running HALC, AbySS [4] version 2.0.2 was used to generate contigs from short reads using 64 as k-mer length as what is suggested in its paper. HALC was run with -o to use ordinary mode so that LoRDEC is used to refine the repeat corrected regions.
- CoLoRMap: OEA mode was not tested in the experiment since it did not improve correction quality significantly but increased its runtime by about two-fold. No other parameters are available for tuning.
- Jabba: version 1.0.0 was used. Prior to running Jabba, karect [5] and btownie (`https://github.com/biointec/brownie`) were run to construct and correct de Bruijn graph with short reads. The de Bruijn k-mer size was set to 75 (-k 75) according to the user-manuals of btownie and Jabba.
- Nanocorr: the long reads were partitioned using the script provided 'partition.py'. The number of reads per file was set to 100, and the number of partition files per directory was also set to 100. The read files in the same directory were processed in parallel. To reduce disk usage, the temporary files were removed immediately after the extraction of corrected reads.
- proovread: version 2.14.1 was used. The coverage of short reads was set according to the data sets used.
- LoRDEC: version 0.8 was built with GATB version 1.41 (`https://github.com/GATB/gatb-core`). Default parameters were used.
- ECTools: the long reads were partitioned and processed in the same way as Nanocorr. AbySS version 2.0.2 was also used to generate contigs from short reads using the same parameters that were used for HALC. Nucmer version 3.1 was used for alignment.
- LSC: version 2.0.1 was used. The default parameters were used.
- FLAS: commit 053c19b was used. The default parameters were used.
- LoRMA: version 0.5 was built with GATB version 1.4.0. LoRDEC version 0.8 was used. Default parameters were used for number of friends (-friends 7) and k-mer size (-k 19).
- Canu: version 1.7 was used. -correct was used to generate corrected reads rather than running the whole assembly pipeline. The genome size was set to the size of the reference genome. Parameter 'useGrid' was set to false to run Canu on one node. Parameter 'stopOnReadQuality' was also set to false to keep Canu running even if read quality is low. Parameters '-pacbio-raw' and '-nanopore-raw' were used for PacBio and ONT reads respectively.

Table S1: Subsamples of D2-P and D2-O

| Data set | Sequencing depth | N50 (bp) | Number of reads |
|---|---|---|---|
| D2-P-10x | 10x | 8635 | 20113 |
| D2-P-20x | 20x | 8665 | 39989 |
| D2-P-30x | 30x | 8674 | 59928 |
| D2-P-60x | 60x | 8646 | 119717 |
| D2-P-90x | 90x | 8649 | 179488 |
| D2-O-10x | 10x | 6976 | 20449 |
| D2-O-20x | 20x | 6983 | 40720 |
| D2-O-30x | 30x | 6994 | 61292 |

Table S2: Experimental results for D2-P-10x

| Method | # Reads | # Bases (Mbp) | # Aligned reads | # Aligned bases (Mbp) | Maximum length (bp) | N50 (bp) | Genome fraction (%) | Alignment identity (%) | CPU time (hh:mm:ss) | Wall time (hh:mm:ss) | Memory Usage (GB) |
|---|---|---|---|---|---|---|---|---|---|---|---|
| Original | 20 113 | 122.6 | 19 801 | 111.9 | 28 392 | 8635 | 99.953 | 87.2209 | - | - | - |
| *Non-hybrid methods* | | | | | | | | | | | |
| FLAS | 15 474 | 79.8 | 15 471 | 78.5 | 26 236 | 6363 | 90.204 | 97.4409 | 00:14:53 | 00:02:39 | 1.1 |
| LoRMA | 9641 | 11.0 | 9626 | 10.9 | 9370 | 1260 | 3.775 | 99.3906 | 02:11:54 | 00:08:51 | 63.7 |
| Canu | 18 747 | 113.4 | 18 614 | 105.2 | 27 582 | 8258 | 99.919 | 95.6321 | 01:43:19 | 00:07:16 | 1.7 |
| *Short-read-assembly-based methods* | | | | | | | | | | | |
| HG-CoLoR | 19 945 | 114.6 | 19 933 | 109.8 | 30 409 | 8108 | 99.971 | 99.3121 | 41:07:30 | 02:00:39 | 30.8 |
| FMLRC | 20 056 | 115.8 | 19 901 | 110.0 | 27 851 | 8149 | 99.972 | 99.3699 | 00:28:13 | 00:12:16 | 5.5 |
| HALC | 20 061 | 117.0 | 20 000 | 108.2 | 28 792 | 8231 | 99.971 | 99.0492 | 06:04:58 | 00:22:16 | 3.9 |
| Jabba | 16 994 | 90.7 | 16 987 | 90.7 | 28 392 | 7831 | 94.619 | 99.9825 | 00:27:21 | 00:03:49 | 21.4 |
| LoRDEC | 20 063 | 117.8 | 19 936 | 109.0 | 27 963 | 8296 | 99.968 | 97.9255 | 01:59:16 | 00:07:25 | 1.6 |
| ECTools | 11 029 | 79.6 | 11 025 | 79.3 | 27 593 | 8390 | 99.103 | 99.7681 | 56:00:03 | 03:31:58 | 4.3 |
| *Short-read-alignment-based methods* | | | | | | | | | | | |
| Hercules | - | - | - | - | - | - | - | - | - | >72:00:00 | - |
| CoLoRMap | 20 110 | 120.2 | 19 984 | 112.3 | 28 299 | 8331 | 99.973 | 97.7314 | 09:05:14 | 00:35:53 | 17.6 |
| Nanocorr | 19 462 | 106.8 | 19 455 | 104.4 | 28 515 | 7885 | 99.944 | 98.8748 | 375:05:59 | 15:54:35 | 23.3 |
| proovread | 20 053 | 117.4 | 19 935 | 109.4 | 28 257 | 8270 | 99.965 | 98.2908 | 13:50:11 | 01:21:28 | 14.6 |
| LSC | 19 310 | 110.0 | 19 284 | 104.6 | 27 642 | 8099 | 99.953 | 96.2482 | 415:34:35 | 19:22:04 | 82.6 |

Table S3: Experimental results for D2-P-20x

| Method | # Reads | # Bases (Mbp) | # Aligned reads | # Aligned bases (Mbp) | Maximum length (bp) | N50 (bp) | Genome fraction (%) | Alignment identity (%) | CPU time (hh:mm:ss) | Wall time (hh:mm:ss) | Memory Usage (GB) |
|---|---|---|---|---|---|---|---|---|---|---|---|
| Original | 39 989 | 244.1 | 39 358 | 223.0 | 31 859 | 8665 | 99.976 | 87.2611 | - | - | - |
| *Non-hybrid methods* | | | | | | | | | | | |
| FLAS | 30 822 | 179.5 | 30 807 | 177.2 | 27 839 | 7383 | 99.753 | 98.6296 | 00:40:04 | 00:06:01 | 2.2 |
| LoRMA | 18 827 | 19.4 | 18 798 | 19.3 | 10 847 | 1095 | 5.388 | 99.2904 | 07:31:50 | 00:24:43 | 63.9 |
| Canu | 37 712 | 227.6 | 37 404 | 211.0 | 31 058 | 8279 | 99.974 | 97.0266 | 04:11:45 | 00:14:34 | 4.7 |
| *Short-read-assembly-based methods* | | | | | | | | | | | |
| HG-CoLoR | - | - | - | - | - | - | - | - | - | - | - |
| FMLRC | 39 879 | 230.6 | 39 574 | 219.3 | 30 884 | 8192 | 99.976 | 99.3929 | 01:12:06 | 00:14:38 | 5.5 |
| HALC | 39 893 | 233.0 | 39 777 | 222.1 | 30 871 | 8283 | 99.976 | 98.9419 | 09:43:42 | 01:16:27 | 25.5 |
| Jabba | 33 709 | 181.2 | 33 694 | 181.1 | 30 110 | 7934 | 94.973 | 99.9843 | 00:27:23 | 00:03:46 | 21.4 |
| LoRDEC | 39 895 | 234.6 | 39 630 | 217.0 | 30 871 | 8336 | 99.976 | 97.9517 | 04:19:37 | 00:13:12 | 1.6 |
| ECTools | 21 778 | 158.5 | 21 771 | 157.9 | 27 593 | 8457 | 99.483 | 99.7685 | 115:09:51 | 06:48:31 | 4.3 |
| *Short-read-alignment-based methods* | | | | | | | | | | | |
| Hercules | - | - | - | - | - | - | - | - | - | >72:00:00 | - |
| CoLoRMap | 39 979 | 239.3 | 39 696 | 222.8 | 30 860 | 8398 | 99.975 | 97.5788 | 08:54:06 | 00:48:47 | 20.6 |
| Nanocorr | 37 081 | 204.0 | 37 065 | 199.3 | 28 515 | 7928 | 99.974 | 98.8683 | 748:27:25 | 28:25:25 | 23.9 |
| proovread | 39 894 | 234.0 | 39 636 | 217.5 | 30 818 | 8319 | 99.977 | 98.1525 | 21:39:23 | 02:31:41 | 47.9 |
| LSC | 28 386 | 162.9 | 28 361 | 155.7 | 28 097 | 8185 | 99.975 | 96.4823 | 900:15:49 | 37:05:17 | 101.9 |

Note: HG-CoLoR reported an error when correcting this dataset.

Table S4: Experimental results for D2-P-30x

| Method | # Reads | # Bases (Mbp) | # Aligned reads | # Aligned bases (Mbp) | Maximum length (bp) | N50 (bp) | Genome fraction (%) | Alignment identity (%) | CPU time (hh:mm:ss) | Wall time (hh:mm:ss) | Memory Usage (GB) |
|---|---|---|---|---|---|---|---|---|---|---|---|
| Original | 59 928 | 366.1 | 58 969 | 334.1 | 31 859 | 8674 | 99.976 | 87.2517 | - | - | - |
| *Non-hybrid methods* | | | | | | | | | | | |
| FLAS | 46 430 | 281.2 | 46 407 | 277.5 | 30 823 | 7779 | 99.972 | 99.0321 | 01:20:05 | 00:10:32 | 2.5 |
| LoRMA | 36 267 | 34.5 | 36 238 | 34.3 | 9973 | 976 | 9.614 | 99.3186 | 15:39:16 | 00:44:10 | 64.2 |
| Canu | 48 936 | 283.1 | 48 906 | 279.5 | 28 146 | 7985 | 99.974 | 99.1508 | 06:44:09 | 00:22:21 | 6.3 |
| *Short-read-assembly-based methods* | | | | | | | | | | | |
| HG-CoLoR | 59 071 | 340.5 | 59 031 | 325.9 | 45 830 | 8129 | 99.975 | 99.0657 | 109:40:58 | 05:00:33 | 52.5 |
| FMLRC | 59 762 | 345.7 | 59 288 | 328.6 | 30 884 | 8192 | 99.976 | 99.3896 | 02:05:00 | 00:15:03 | 5.5 |
| HALC | 59 784 | 349.3 | 59 598 | 322.8 | 30 864 | 8285 | 99.976 | 99.0676 | 13:38:33 | 02:17:41 | 21.7 |
| Jabba | 50 651 | 271.9 | 50 628 | 271.8 | 30 110 | 7897 | 95.104 | 99.9834 | 00:30:24 | 00:03:57 | 21.4 |
| LoRDEC | 59 795 | 351.7 | 59 392 | 325.2 | 30 871 | 8335 | 99.976 | 97.9507 | 06:29:45 | 00:18:34 | 1.6 |
| ECTools | 32 701 | 237.5 | 32 689 | 236.5 | 28 107 | 8437 | 99.598 | 99.7714 | 178:10:23 | 10:50:32 | 4.5 |
| *Short-read-alignment-based methods* | | | | | | | | | | | |
| Hercules | - | - | - | - | - | - | - | - | - | >72:00:00 | - |
| CoLoRMap | 59 911 | 358.2 | 59 421 | 332.4 | 31 129 | 8397 | 99.976 | 97.1147 | 09:15:34 | 01:00:21 | 22.0 |
| Nanocorr | 56 946 | 313.2 | 56 922 | 306.1 | 28 515 | 7901 | 99.974 | 98.8705 | 1119:49:00 | 42:22:05 | 24.4 |
| proovread | 59 794 | 351.3 | 59 364 | 325.8 | 30 820 | 8327 | 99.976 | 97.9828 | 31:41:40 | 03:51:31 | 19.5 |
| LSC | 57 501 | 328.8 | 57 430 | 312.3 | 31 242 | 8156 | 99.976 | 96.2465 | 1503:43:21 | 56:50:09 | 126.9 |

Table S5: Experimental results for D2-P-60x

| Method | # Reads | # Bases (Mbp) | # Aligned reads | # Aligned bases (Mbp) | Maximum length (bp) | N50 (bp) | Genome fraction (%) | Alignment identity (%) | CPU time (hh:mm:ss) | Wall time (hh:mm:ss) | Memory Usage (GB) |
|---|---|---|---|---|---|---|---|---|---|---|---|
| Original | 119 717 | 731.0 | 117 797 | 666.0 | 33 569 | 8646 | 99.976 | 87.2680 | - | - | - |
| *Non-hybrid methods* | | | | | | | | | | | |
| FLAS | 90 214 | 560.0 | 90 154 | 552.3 | 30 926 | 8019 | 99.975 | 99.4142 | 4:08:56 | 0:28:08 | 4.6 |
| LoRMA | 327 985 | 413.8 | 327 922 | 413.5 | 14 691 | 1495 | 82.967 | 99.7697 | 53:25:06 | 02:21:48 | 66.6 |
| Canu | 62 265 | 322.5 | 62 236 | 319.4 | 27 767 | 7280 | 99.940 | 99.2633 | 05:37:49 | 00:19:42 | 6.3 |
| *Short-read-assembly-based methods* | | | | | | | | | | | |
| HG-CoLoR | - | - | - | - | - | - | - | - | - | - | - |
| FMLRC | 119 359 | 690.1 | 118 424 | 655.5 | 33 643 | 8178 | 99.976 | 99.3838 | 03:50:30 | 00:19:47 | 5.4 |
| HALC | 119 393 | 697.4 | 119 041 | 643.7 | 33 391 | 8266 | 99.976 | 99.0768 | 27:42:21 | 04:31:29 | 22.3 |
| Jabba | 101 451 | 544.2 | 101 405 | 543.9 | 30 110 | 7865 | 95.380 | 99.9832 | 00:38:22 | 00:04:21 | 21.4 |
| LoRDEC | 119 425 | 702.2 | 118 623 | 648.5 | 33 414 | 8318 | 99.976 | 97.9590 | 12:55:31 | 00:31:52 | 1.7 |
| ECTools | 65 492 | 473.9 | 65 474 | 472.0 | 28 186 | 8414 | 99.705 | 99.7703 | 386:47:01 | 22:13:17 | 4.4 |
| *Short-read-alignment-based methods* | | | | | | | | | | | |
| Hercules | 119 647 | 724.3 | 117 814 | 660.0 | 33 533 | 8575 | 99.976 | 90.1448 | 364:22:49 | 13:13:01 | 247.7 |
| CoLoRMap | 119 668 | 714.6 | 118 558 | 660.5 | 33 398 | 8401 | 99.976 | 96.5449 | 11:52:58 | 01:38:33 | 25.4 |
| Nanocorr | - | - | - | - | - | - | - | - | - | >72:00:00 | - |
| proovread | 119 457 | 702.9 | 118 424 | 649.1 | 33 341 | 8333 | 99.976 | 97.4479 | 71:11:36 | 08:18:46 | 29.9 |
| LSC | - | - | - | - | - | - | - | - | - | >72:00:00 | - |

Note: HG-CoLoR reported an error when correcting this dataset.

Table S6: Experimental results for D2-P-90x

| Method | # Reads | # Bases (Mbp) | # Aligned reads | # Aligned bases (Mbp) | Maximum length (bp) | N50 (bp) | Genome fraction (%) | Alignment identity (%) | CPU time (hh:mm:ss) | Wall time (hh:mm:ss) | Memory Usage (GB) |
|---|---|---|---|---|---|---|---|---|---|---|---|
| Original | 179 488 | 1095.9 | 176 591 | 998.2 | 35 196 | 8649 | 99.976 | 87.2655 | - | - | - |
| *Non-hybrid methods* | | | | | | | | | | | |
| FLAS | 132 435 | 829.5 | 132 330 | 818.4 | 30 200 | 8076 | 99.976 | 99.5446 | 07:52:02 | 00:50:35 | 6.3 |
| LoRMA | 473 099 | 848.4 | 472 994 | 848.0 | 19 426 | 2405 | 99.589 | 99.8003 | 112:39:38 | 04:42:55 | 69.3 |
| Canu | 64 459 | 459.1 | 64 440 | 454.3 | 28 521 | 8190 | 99.976 | 99.4384 | 09:13:31 | 00:30:03 | 6.7 |
| *Short-read-assembly-based methods* | | | | | | | | | | | |
| HG-CoLoR | - | - | - | - | - | - | - | - | - | - | - |
| FMLRC | 178 954 | 1034.5 | 177 577 | 982.3 | 33 658 | 8180 | 99.976 | 99.3872 | 05:54:38 | 00:24:10 | 5.5 |
| HALC | 179 016 | 1045.5 | 178 487 | 964.7 | 34 783 | 8262 | 99.976 | 99.0738 | 39:00:28 | 06:40:52 | 27.3 |
| Jabba | 152 100 | 814.6 | 152 021 | 814.2 | 30 141 | 7849 | 95.514 | 99.9831 | 00:34:37 | 00:04:36 | 21.4 |
| LoRDEC | 179 063 | 1052.7 | 177 874 | 971.8 | 34 896 | 8321 | 99.978 | 97.9513 | 19:51:52 | 00:46:31 | 1.9 |
| ECTools | 98 063 | 709.0 | 98 036 | 706.2 | 28 749 | 8408 | 99.728 | 99.7721 | 643:09:22 | 42:43:28 | 4.6 |
| *Short-read-alignment-based methods* | | | | | | | | | | | |
| Hercules | 179 444 | 1091.5 | 176 614 | 994.3 | 35 196 | 8619 | 99.976 | 88.4041 | 205:47:53 | 07:31:11 | 248.0 |
| CoLoRMap | 179 406 | 1071.1 | 177 772 | 990.0 | 35 107 | 8403 | 99.976 | 96.4924 | 15:29:41 | 02:21:01 | 30.1 |
| Nanocorr | - | - | - | - | - | - | - | - | - | >72:00:00 | - |
| proovread | 179 133 | 1056.1 | 177 376 | 972.2 | 35 088 | 8347 | 96.814 | 96.8140 | 120:38:36 | 12:43:12 | 39.1 |
| LSC | - | - | - | - | - | - | - | - | - | >72:00:00 | - |

Note: HG-CoLoR reported an error when correcting this dataset.

Table S7: Experimental results for subsamples of D2-O-10x

| Method | # Reads | # Bases (Mbp) | # Aligned reads | # Aligned bases (Mbp) | Maximum length (bp) | N50 (bp) | Genome fraction (%) | Alignment identity (%) | CPU time (hh:mm:ss) | Wall time (hh:mm:ss) | Memory Usage (GB) |
|---|---|---|---|---|---|---|---|---|---|---|---|
| Original | 20 225 | 121.6 | 18 472 | 108.4 | 55 374 | 6979 | 99.896 | 86.1285 | - | - | - |
| *Non-hybrid methods* | | | | | | | | | | | |
| FLAS | 16 528 | 82.5 | 16 476 | 81.8 | 17 924 | 5623 | 91.721 | 94.7627 | 00:27:46 | 00:03:12 | 0.3 |
| LoRMA | 10 650 | 14.2 | 10 488 | 13.9 | 9155 | 1653 | 6.391 | 97.8597 | 02:31:10 | 00:09:17 | 63.7 |
| Canu | 19 236 | 115.5 | 18 179 | 106.5 | 20 355 | 6899 | 99.631 | 94.6333 | 02:04:50 | 00:08:33 | 5.7 |
| *Short-read-assembly-based methods* | | | | | | | | | | | |
| HG-CoLoR | - | - | - | - | - | - | - | - | - | - | - |
| FMLRC | 20 219 | 121.3 | 19 010 | 113.3 | 55 374 | 6979 | 99.947 | 99.2462 | 00:47:07 | 00:11:15 | 2.2 |
| HALC | 20 218 | 122.0 | 19 451 | 110.0 | 55 377 | 7011 | 99.939 | 98.8731 | 05:57:43 | 00:22:14 | 4.0 |
| Jabba | 16 782 | 91.1 | 16 723 | 91.0 | 20 379 | 6726 | 94.506 | 99.9806 | 00:36:33 | 00:03:37 | 21.5 |
| LoRDEC | 20 227 | 122.5 | 18 836 | 110.1 | 55 375 | 7029 | 99.932 | 96.8841 | 01:42:07 | 00:08:02 | 1.7 |
| ECTools | 13 817 | 90.4 | 13 774 | 89.9 | 20 358 | 7053 | 98.658 | 99.7718 | 74:06:02 | 03:46:36 | 5.3 |
| *Short-read-alignment-based methods* | | | | | | | | | | | |
| Hercules | - | - | - | - | - | - | - | - | - | >72:00:00 | - |
| CoLoRMap | 20 247 | 123.3 | 18 684 | 111.7 | 55 374 | 7051 | 99.936 | 97.3563 | 06:49:34 | 00:30:13 | 13.4 |
| Nanocorr | 17 728 | 103.0 | 17 675 | 102.2 | 20 532 | 6929 | 99.879 | 98.7603 | 631:40:01 | 24:23:08 | 14.5 |
| proovread | 20 223 | 121.8 | 18 693 | 109.9 | 55 374 | 6994 | 99.942 | 97.6638 | 11:52:55 | 01:11:43 | 14.1 |
| LSC | 17 103 | 99.1 | 17 041 | 98.1 | 20 452 | 6806 | 99.870 | 94.8926 | 676:27:14 | 27:08:41 | 147.8 |

Note: HG-CoLoR reported an error when correcting this dataset.

Table S8: Experimental results for subsamples of D2-O-20x

| Method | # Reads | # Bases (Mbp) | # Aligned reads | # Aligned bases (Mbp) | Maximum length (bp) | N50 (bp) | Genome fraction (%) | Alignment identity (%) | CPU time (hh:mm:ss) | Wall time (hh:mm:ss) | Memory Usage (GB) |
|---|---|---|---|---|---|---|---|---|---|---|---|
| Original | 40 299 | 242.4 | 36 821 | 215.8 | 55 374 | 6985 | 99.966 | 86.1341 | - | - | - |
| *Non-hybrid methods* | | | | | | | | | | | |
| FLAS | 33 868 | 193.7 | 33 763 | 192.3 | 21 785 | 6374 | 99.660 | 96.1600 | 01:19:51 | 00:07:54 | 1.9 |
| LoRMA | 52 979 | 51.4 | 52 785 | 50.9 | 9147 | 997 | 35.481 | 98.2431 | 09:07:28 | 00:29:41 | 64.2 |
| Canu | 38 609 | 232.3 | 36 309 | 213.8 | 42 911 | 6913 | 99.856 | 95.6186 | 04:59:32 | 00:14:45 | 7.2 |
| *Short-read-assembly-based methods* | | | | | | | | | | | |
| HG-CoLoR | - | - | - | - | - | - | - | - | - | - | - |
| FMLRC | 40 288 | 241.8 | 37 931 | 225.6 | 55 374 | 6979 | 99.967 | 99.2302 | 01:21:36 | 00:12:19 | 1.3 |
| HALC | 40 293 | 243.3 | 38 719 | 219.2 | 55 378 | 7011 | 99.972 | 98.8497 | 16:35:41 | 01:26:22 | 4.1 |
| Jabba | 33 467 | 181.4 | 33 333 | 181.1 | 20 379 | 6727 | 94.950 | 99.9825 | 00:39:48 | 00:03:46 | 21.5 |
| LoRDEC | 40 301 | 244.2 | 37 535 | 219.2 | 55 375 | 7033 | 99.968 | 96.8777 | 03:33:04 | 00:12:56 | 1.8 |
| ECTools | 27 436 | 179.6 | 27 354 | 178.7 | 20 358 | 7059 | 99.080 | 99.7703 | 144:19:40 | 06:48:58 | 5.6 |
| *Short-read-alignment-based methods* | | | | | | | | | | | |
| Hercules | - | - | - | - | - | - | - | - | - | >72:00:00 | - |
| CoLoRMap | 40 333 | 245.6 | 37 127 | 221.8 | 55 374 | 7054 | 99.971 | 97.0285 | 07:36:44 | 00:45:04 | 20.7 |
| Nanocorr | 35 299 | 205.0 | 35 187 | 203.4 | 23 384 | 6929 | 99.965 | 98.7633 | 1260:36:46 | 47:38:41 | 15.2 |
| proovread | 40 297 | 242.9 | 37 190 | 218.7 | 55 374 | 6995 | 99.971 | 97.3920 | 20:03:49 | 02:15:13 | 15.9 |
| LSC | 34 067 | 197.5 | 33 943 | 195.4 | 23 499 | 6810 | 99.959 | 94.9089 | 1323:52:37 | 49:05:31 | 106.5 |

Note: HG-CoLoR reported an error when correcting this dataset.

Table S9: Experimental results for subsamples of D2-O-30x

| Method | # Reads | # Bases (Mbp) | # Aligned reads | # Aligned bases (Mbp) | Maximum length (bp) | N50 (bp) | Genome fraction (%) | Alignment identity (%) | CPU time (hh:mm:ss) | Wall time (hh:mm:ss) | Memory Usage (GB) |
|---|---|---|---|---|---|---|---|---|---|---|---|
| Original | 60 648 | 365.4 | 55 402 | 325.4 | 55 374 | 6997 | 99.971 | 86.1812 | - | - | - |
| *Non-hybrid methods* | | | | | | | | | | | |
| FLAS | 50 245 | 300.1 | 50 085 | 298.0 | 21 411 | 6686 | 99.909 | 96.5553 | 02:30:09 | 00:13:51 | 2.7 |
| LoRMA | 154 864 | 146.3 | 154 668 | 145.7 | 9512 | 972 | 78.164 | 98.4020 | 20:18:16 | 00:59:45 | 65.3 |
| Canu | 49 872 | 302.3 | 49 748 | 300.2 | 23 194 | 6966 | 99.880 | 97.2080 | 08:40:23 | 00:26:48 | 6.7 |
| *Short-read-assembly-based methods* | | | | | | | | | | | |
| HG-CoLoR | - | - | - | - | - | - | - | - | - | - | - |
| FMLRC | 60 635 | 364.5 | 57 134 | 340.2 | 55 374 | 6990 | 99.970 | 99.2438 | 01:56:19 | 00:13:31 | 2.2 |
| HALC | 60 644 | 366.8 | 58 310 | 330.6 | 55 377 | 7023 | 99.972 | 98.8575 | 23:56:13 | 01:43:55 | 4.1 |
| Jabba | 50 474 | 273.4 | 50 271 | 272.9 | 20 379 | 6732 | 95.124 | 99.9816 | 00:44:06 | 00:03:59 | 21.5 |
| LoRDEC | 60 652 | 368.1 | 56 507 | 330.5 | 55 375 | 7047 | 99.971 | 96.9152 | 05:18:57 | 00:15:32 | 1.9 |
| ECTools | 41 303 | 270.8 | 41 182 | 269.5 | 20 358 | 7073 | 99.174 | 99.7668 | 216:17:19 | 10:20:44 | 5.7 |
| *Short-read-alignment-based methods* | | | | | | | | | | | |
| Hercules | - | - | - | - | - | - | - | - | - | >72:00:00 | - |
| CoLoRMap | 60 693 | 369.5 | 55 812 | 333.0 | 55 374 | 7055 | 99.972 | 96.5312 | 08:30:17 | 00:58:56 | 22.9 |
| Nanocorr | 53 159 | 309.4 | 52 994 | 306.9 | 23 384 | 6947 | 99.971 | 98.7687 | 1898:17:48 | 71:13:41 | 15.0 |
| proovread | 60 649 | 366.2 | 55 910 | 329.4 | 55 374 | 7009 | 99.974 | 97.2014 | 30:07:31 | 03:25:32 | 19.4 |
| LSC | - | - | - | - | - | - | - | - | - | >72:00:00 | - |

Note: HG-CoLoR reported an error when correcting this dataset.

### References


[1] Ben Langmead and Steven L Salzberg. Fast gapped-read alignment with bowtie 2. *Nature methods*, 9(4):357, 2012.

[2] Mark J Chaisson and Glenn Tesler. Mapping single molecule sequencing reads using basic local alignment with successive refinement (blasr): application and theory. *BMC bioinformatics*, 13(1):238, 2012.

[3] James Holt and Leonard McMillan. Merging of multi-string bwts with applications. *Bioinformatics*, 30(24):3524–3531, 2014.

[4] Shaun D Jackman, Benjamin P Vandervalk, Hamid Mohamadi, Justin Chu, Sarah Yeo, S Austin Hammond, Golnaz Jahesh, Hamza Khan, Lauren Coombe, Rene L Warren, et al. Abyss 2.0: resource-efficient assembly of large genomes using a bloom filter. *Genome research*, 27(5):768–777, 2017.

[5] Amin Allam, Panos Kalnis, and Victor Solovyev. Karect: accurate correction of substitution, insertion and deletion errors for next-generation sequencing data. *Bioinformatics*, 31(21):3421–3428, 2015.
